# Genetic dissection of Nodal and Bmp signalling requirements during primordial germ cell development

**DOI:** 10.1101/464776

**Authors:** Anna D. Senft, Elizabeth K. Bikoff, Elizabeth J. Robertson, Ita Costello

## Abstract

The essential roles played by Nodal and Bmp signalling during early mouse development have been extensively documented. Here we used conditional deletion strategies to investigate functional contributions made by Nodal, Bmp and Smad downstream effectors during primordial germ cell (PGC) development. We demonstrate that Nodal and its target gene Eomes provide early instructions during formation of the PGC lineage. We discovered that Smad2 inactivation in the visceral endoderm results in increased numbers of PGCs due to an expansion of the PGC niche. Smad1 is required for specification, whereas in contrast Smad4 controls the maintenance and migration of PGCs. Importantly we found that beside Blimp1, down-regulated phosphoSmad159 levels also distinguishes PGCs from their somatic neighbours so that emerging PGCs become refractory to Bmp signalling that otherwise promotes mesodermal development in the posterior epiblast. Thus balanced Nodal/Bmp signalling cues regulate germ cell versus somatic cell fate decisions in the early posterior epiblast.

## Introduction

Primordial germ cells (PGCs), the precursors of sperm and eggs, are initially detectable in the early mouse embryo at around embryonic day (e) 6.25, prior to the onset of gastrulation^1^. Early fate mapping experiments revealed that the proximal posterior epiblast (PPE) gives rise to both the extra-embryonic mesoderm (ExM) and PGC cell populations^2^. The regulatory signals governing these cell fate decisions remain ill-defined. The PR domain containing zinc finger transcription factor Blimp1 (encoded by *Prdm1*) plays an essential role during PGC specification and expression at e6.25 identifies precursor PGCs (pre-PGCs)^3,4^. Commitment to the PGC lineage becomes evident slightly later between e6.75-e7.5 when pre-PGCs activate expression of germ cell markers such as Stella (*Dppa3*) and Ap2γ (*Tfap2c*), concomitantly reactivate expression of pluripotency genes including Sox2 and repress somatic gene expression^1,5^. By e8.5, specified PGCs have migrated from the base of the allantois into the overlying endoderm, and subsequently migrate along the dorsal hindgut endoderm before homing to and colonising the genital ridges from e10.5^1^. Extensive chromatin remodelling and genome-wide epigenetic reprogramming occurs during migration^6^. Transcriptional profiling experiments analysing PGCs *in vivo* and *in vitro* differentiated PGC-like cells (PGCLCs) have provided new insights into the dynamic transcriptional changes that accompany PGC maturation^5,7,8^.

Correct patterning of the early post-implantation stage embryo depends on reciprocal signalling cues by members of the TGFβ family of secreted growth factors controlling cell-cell interactions between the pluripotent epiblast and the overlying extra-embryonic tissues namely the extra-embryonic ectoderm (ExE) and the visceral endoderm (VE)^9,10^. Temporally and spatially restricted expression of Nodal and Bmp ligands results in activation of their cognate receptors, phosphorylation of the intracellular effectors, Smad2 and Smad3 (Smad23), or Smad1, Smad5 and Smad9 (Smad159) respectively, that associate with the co-Smad Smad4, translocate to the nucleus and activate cell-type specific target gene expression^11^. Nodal signalling in the early epiblast is required for correct patterning of the VE, formation of the anterior-posterior (A-P) axis and also promotes development of the ExE^12^. Together with Bmp signals from the ExE, Nodal initiates mesoderm induction and primitive streak formation (PS) within the PPE^12,13^. During gastrulation, graded Nodal signals pattern the PS^10^. Thus, highest levels of Nodal signalling are required for specification of anterior PS derivatives. Lowering Nodal expression levels in the PS, or depleting Smad23 in the epiblast results in failure to correctly specify the anterior definitive endoderm (DE) and the embryonic midline^14^.

In contrast Bmp/Smad159 signals promote the formation of ExM. Loss of Bmp4 from the ExE disrupts gastrulation and is associated with truncation of posterior structures including the allantois and yolk sac^13^. Both Smad1 and Smad5 null embryos also display ExM tissue defects^15,16^. Mutant embryos lacking Bmp4 or Bmp8b expression in the ExE or those lacking Bmp2 in the VE display compromised PGC development in the epiblast^17–19^. Bmp4 null embryos entirely lack mature PGCs, while Bmp4 heterozygous embryos contains reduced numbers of PGCs. The observation of reduced numbers of PCGs in embryos lacking either Smad1 or Smad5^15,16^, as well as in Smad1/5 double heterozygous embryos^20^, provides convincing evidence that dose-dependent Bmp signalling governs PGC specification. The Wnt signalling pathway also regulates PGC development^21^. Wnt3 is normally induced in the posterior epiblast in response to Nodal/Bmp signalling^22^. Wnt3 mutants fail to gastrulate^23^ and lack a detectable pre-PGC population^21^. Collectively, these findings demonstrate that patterning during gastrulation and PGC formation are co-ordinately regulated by dynamic signalling events. However, given the complex morphological disturbances observed in loss of function mutant embryos, it has proven difficult to further dissect the crosstalk between the embryonic and extra-embryonic tissues. In particular since Nodal null embryos arrest prior to gastrulation^24^, any possible role of Nodal signalling during PGC specification remains to be explored.

Here we exploited tissue specific conditional deletion strategies to investigate functional contributions made by Nodal and Bmp signalling within the embryonic versus extra-embryonic tissues. We have directly assessed the distinct roles played by various signalling pathway components in the VE and the epiblast during formation of the PGC lineage. Collectively our experiments provide new insights into the signalling cues that cooperatively regulate the size of the founding pre-PGC population and govern the PGC developmental programme during induction, specification and migration at early post-implantation stages of mouse development.

## Results

### Smad2 is required in the VE to restrict Bmp signalling to the proximal region

In Smad2 null embryos, lacking the anterior visceral endoderm (AVE) signalling centre, the epiblast adopts an exclusively ExM fate^12,25^. These mis-patterned embryos arrest at around e9.5. Here, to examine cell type specific Smad2 functional contributions within the VE, we crossed animals carrying a conditional Smad2 allele (Smad2CA) with heterozygous Smad2^+/−^ mice carrying the Ttr-Cre transgene^26^ (Supplemental Fig. S1A). As shown below, we found that the Smad2∆VE embryos phenocopy the Smad2^−/−^ embryos, strengthening the idea that the dramatic tissue disturbances observed in the null embryos predominantly reflect the loss of Smad2 signalling in the AVE.

To assess the possible impact on Bmp signalling, we analysed phospho-Smad1/5/9 (p-Smad159) localisation. At e5.5 p-Smad159 expression is normally restricted to the proximal VE. However, p-Smad159 staining in Smad2∆VE e5.5 embryos is detectable throughout the entire VE, including the distal region (Fig. 1A). Whole-mount *in situ* hybridisation experiments demonstrate that *Bmp4* expression remains unchanged, whereas Smad2∆VE embryos lack *Bmp2* transcripts (Supplemental Fig. 1B). Thus, higher levels of p-Smad159 cannot simply be explained due to increased expression of Bmp ligands.

**Figure 1:**
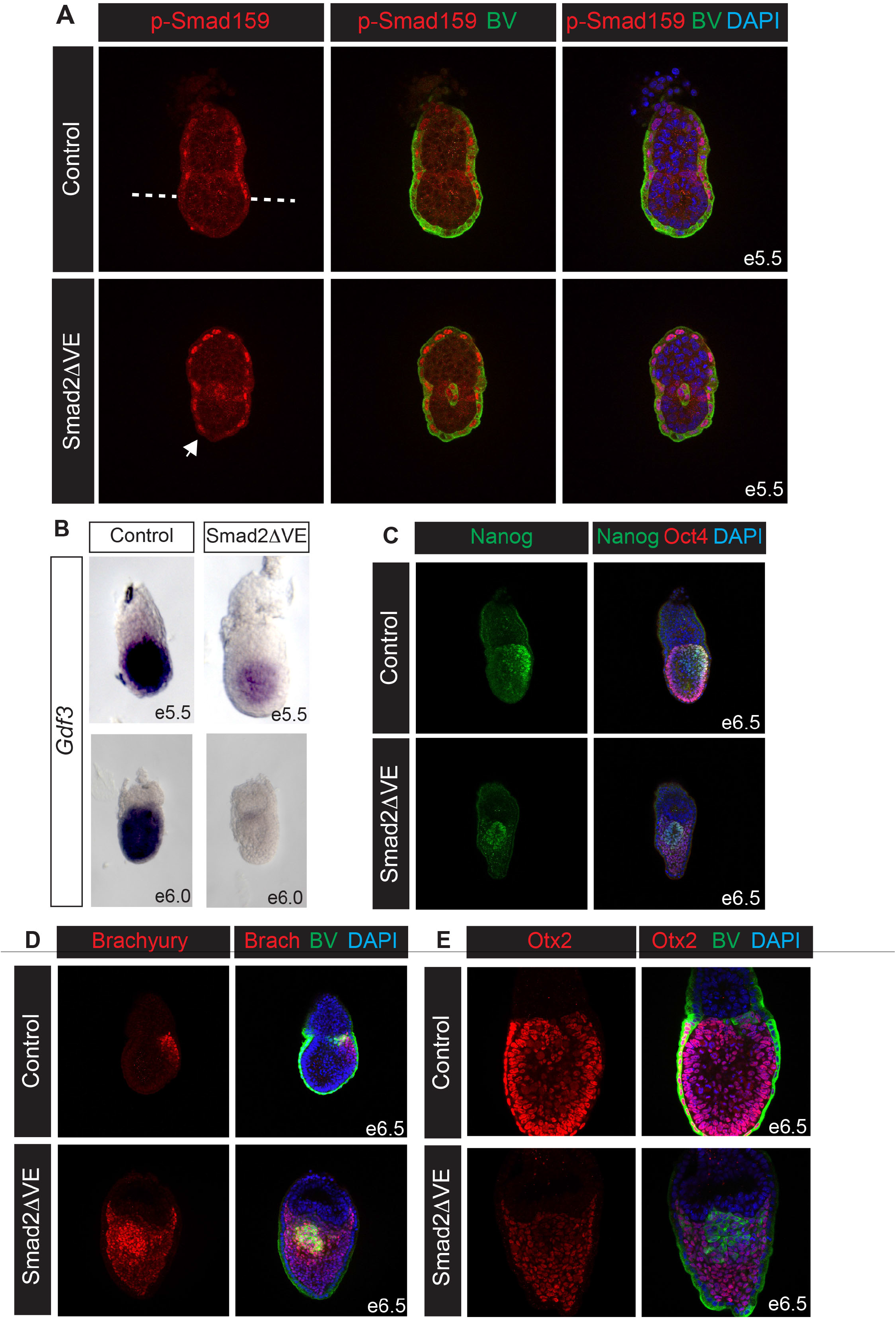
Imbalanced Bmp/Nodal signalling and expansion of the PGC niche caused by Smad2 inactivation in the visceral endoderm. (A) Representative images of p-Smad159 immunofluorescence (IF) staining of e5.5 embryos carrying the Blimp1-Venus (BV) transgene, counterstained with DAPI. Dashed line indicates extent of proximal p-Smad159 staining in control embryos. The arrow indicates expanded p-Smad159 in the distal VE of Smad2∆V embryos. (B) Whole-mount *in-situ* hybridisation analysis of *Gdf3* expression in control and Smad2∆VE embryos at e5.5 and e6.0. (C) Nanog and Oct4 co-staining in e6.5 control and Smad2∆VE embryos. (D) Brachyury IF in e6.5 control and Smad2∆VE BV-expressing embryos. (E). Otx2 staining and BV-expression at e6.5. All IF staining images were counterstained with DAPI.

Antagonistic Bmp and Nodal signalling cues govern VE specification^9^. However, the regulatory mechanisms that normally restrict p-Smad159 signalling to the proximal VE have yet to be fully characterised. The TGFβ antagonist Gdf3, expressed in the epiblast and distal VE, directly antagonises Bmp4 activity^27,28^. Moreover, selective mesoderm expansion in double homozygous embryos lacking both *Gdf3* and the closely related ligand *Gdf1* has been documented^29^. Here we observe in Smad2∆VE embryos that *Gdf3* expression is absent in the VE and reduced in the epiblast (Fig. 1B). Thus, up-regulated p-Smad159 activity in Smad2∆VE embryos potentially reflects decreased *Gdf3* expression levels.

### Conditional inactivation of Smad2 in the visceral endoderm results in expansion of the PGC niche and increased number of PGCs

Alkaline phosphatase (AP) positive PGC clusters were previously identified in e8.5 Smad2^−/−^ embryos^15^. To evaluate PGC formation in Smad2∆VE embryos we examined PGC marker gene expression. Nanog, normally reactivated in the early proximal epiblast^30^, is also strongly expressed in developing PGCs^31^. As expected in control embryos, we detected cells co-expressing Nanog and the pluripotency marker Oct4 in the PPE (Fig. 1C). Similarly, Smad2∆VE e6.5 embryos contain Nanog/Oct4 double positive cells adjacent to the ExE, but these were located in a central position in the epiblast (Fig. 1C).

Next, to assess whether these cells correspond to pre-PGCs, we used the Blimp1-Venus (BV) BAC transgene that faithfully recapitulates Blimp1 expression in both the VE and the developing PGCs^32^. In Smad2DVE mutants expressing the BV transgene we detected BV-positive (BV^+^) pre-PGCs that also co-express E-Cadherin (Supplemental Fig. 1C). As judged by immunohistochemistry these cells strongly express endogenous Blimp1 protein (Supplemental Fig. 1D). Interestingly in Smad2DVE embryos BV^+^ pre-PGCs are initially detectable within the epiblast at e5.5, 12-18 hours before their appearance in wild type embryos (Fig. 1A). Slightly later at e6.5, the Smad2∆VE proximal epiblast contains increased numbers of BV^+^ cells as compared to control wild type embryos (Fig. 1D & 1E).

BV^+^ cells in the proximal epiblast at e6.5 normally express Brachyury and retain E-cadherin expression, whereas the adjacent mesodermal cells also strongly express Brachyury but down-regulate E-cadherin expression (Fig. 1D, Supplemental Fig. 1C). In Smad2∆VE embryos the central core of BV^+^ epithelial cells is likewise surrounded by Brachyury positive mesodermal cells (Fig. 1D). At e6.5, in both control and Smad2∆VE embryos BV^+^ cells weakly express Otx2, Eomes and Sox2 (Fig. 1E, Supplemental Fig. 1E & 1F). Slightly later at e7.5 BV^+^ cell clusters retain ECadherin and co-express Stella as well as Oct4 (Supplemental Fig. 1G & 1H). Overall in Smad2∆VE embryos we observe increased numbers of BV, Oct4, Nanog and Stella co-expressing PGCs surrounded by Brachyury-positive, E-Cadherin-negative mesodermal cells. Thus, Smad2 expression in the VE normally restricts expansion of the PGC niche so that only a small subset of posterior epiblast cells become allocated to a PGC fate.

Next, to examine Smad2 functional contributions within the epiblast we made use of the Sox2-Cre deleter strain^33^. Stella^+^ PGCs are readily detectable at e8.5 in Smad2∆Epi embryos, suggesting that Smad2 is dispensable for PGC specification (Supplemental Fig. 2A). However, it seems likely that the closely related *Smad3* effector, known to be robustly expressed in the epiblast^14^, functionally compensates.

Blimp1 is strongly expressed in the VE but experiments to date have failed to demonstrate any VE-specific functional contributions. To test whether Blimp1 VE expression may influence PGC specification, we used the Ttr-Cre deleter strain. As shown in Supplemental Fig. 2C, selective loss of Blimp1 in the VE has no noticeable effect on the formation of Blimp1 expressing PGCs at the base of the allantois (Supplemental Fig. 2B, 2C). Moreover, Prdm1∆VE mice are viable and fertile (Supplemental Table 1).

### Nodal and its downstream target Eomes play essential roles during PGC specification *in vivo*

Nodal null ES cells, as well as Smad23 double mutant ES, fail to generate PGC-like cells *in* vitro^34,35^. To investigate Nodal functional contributions *in vivo* we examined Nanog expression in Nodal^−/−^ embryos. At e5.5 Nanog expression levels are dramatically reduced as compared to control embryos (Fig. 2A, Supplemental Fig. 2D). However, Nodal^−/−^ embryos robustly express the epiblast marker Oct4 (Supplemental Fig. 2D). Interestingly, Nodal^−/−^ embryos display precocious induction of BV^+^ epiblast cells (Figure 2A), which only weakly express Nanog (Fig. 2A). As for Smad2∆VE embryos, e5.5 Nodal^−/−^ mutants similarly exhibit increased levels of p-Smad159 staining in the distal VE (Fig. 2B). Moreover p-Smad159 is also detectable throughout the epiblast (Fig. 2B), strongly suggesting that Nodal functions *in vivo* to promote optimal levels of Nanog during pre-PGC development and also to spatially restrict the PGC niche.

**Figure 2:**
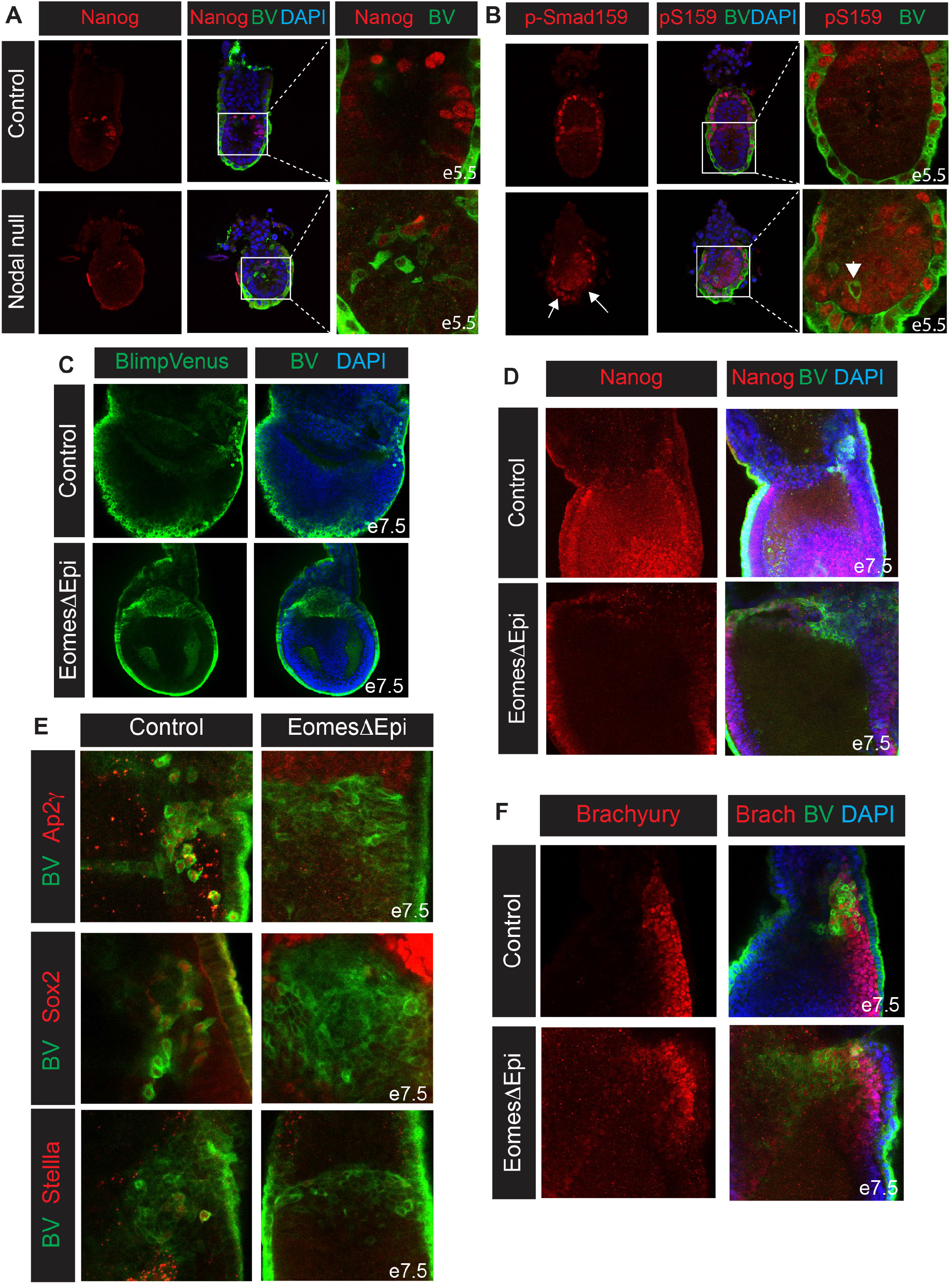
Nodal and its downstream target Eomes regulate early stages of PGC development. (A) Reduced levels of Nanog expression in e5.5 Nodal null BV^+^ embryos. (B) Nodal null BV^+^ embryos display an expansion of p-Smad159 staining in the distal VE, as indicated by arrows. Arrowhead indicates lowered p-Smad159 staining within BV^+^ cells. (C) Expansion of the BV^+^ cell population in EomesΔEpi embryos at e7.5. (D) Nuclear Nanog staining in EomesΔEpi and control BV^+^ embryos at e7.5. (E) Analysis of Ap2γ, Sox2 and Stella in e7.5 control and EomesΔEpi BV^+^ cells. (F) Brachyury staining in control and EomesDEpi BV^+^ e7.5 embryos. All IF images are counterstained with DAPI.

The T-box transcription factor Eomes, acting downstream of Nodal signalling, plays essential roles during gastrulation^36,37^. Conditional deletion in the epiblast (Eomes∆Epi) results in defective epithelial to mesenchymal transition (EMT) and the failure of nascent mesoderm cells to down-regulate E-cadherin and exit the PS^36^. Eomes is expressed in early pre-PGCs but becomes down-regulated by late streak stages (Supplemental Fig. 1E, Supplemental Fig. 2E). Interestingly, Eomes∆Epi e7.5 embryos contain an expanded epithelial-like BV^+^ cell population (Fig. 2C) that also express endogenous Blimp1 protein (Supplemental Fig. 2F). Nanog expression is reactivated within the epiblast, however the BV^+^ cell population in EomesDEpi embryos is largely Nanog negative (Fig. 2D). Moreover, at e7.5 BV^+^ cells fail to activate the PGC markers Ap2γ, Stella and Sox2 (Fig. 2E) and only a subset retain Brachyury activity (Fig. 2F). These results demonstrate that the loss of the T-box transcription factor Eomes in the epiblast disrupts the PGC developmental programme *in vivo*.

Eomes is also required to pattern the VE^38^. As shown in Supplemental Fig. 2F, Eomes∆VE embryos contain Blimp1^+^ epiblast cells. Thus, Eomes activity in the VE is non-essential for Blimp1 induction in the epiblast.

### Smad1 is required for specification whereas Smad4 controls the maintenance and migration of PGCs

Smad4 in association with the receptor Smads, Smad159 activates Bmp target gene expression. As judged by AP staining, both Smad1 null and Smad4∆Epi embryos lack PGCs at e8.5^15,39^. However, cooperative or possibly unique functional roles at distinct developmental stages during PGC lineage specification have yet to be examined. Initially to explore Smad4 functional requirements in the VE, we generated Smad4∆VE embryos. We found at e7.5 that Blimp1-expressing PGCs are formed appropriately at the base of the allantois (Supplemental Fig. 3A). Similarly, Smad4 activity in the epiblast is non-essential for PGC specification. BV/Stella co-expressing cells are detectable on the posterior side of Smad4∆Epi embryos at e7.5 (Supplemental Fig. 3B). However, at e8.5 these Stella/ BV^+^ PGCs remain at the base of the allantois and fail to migrate towards the gut endoderm (Fig. 3A). Additionally, we used a Blimp1-Cre deleter strain^40^ to selectively eliminate Smad4 expression within the pre-PGC cell population (Smad4∆PGC). Relatively few Stella expressing cells were present in e8.5 Smad4∆PGC embryos (Supplemental Fig. 3C). Thus, Smad4 is dispensable for initial PGC specification but is required for PGC maintenance and/or migration.

**Figure 3:**
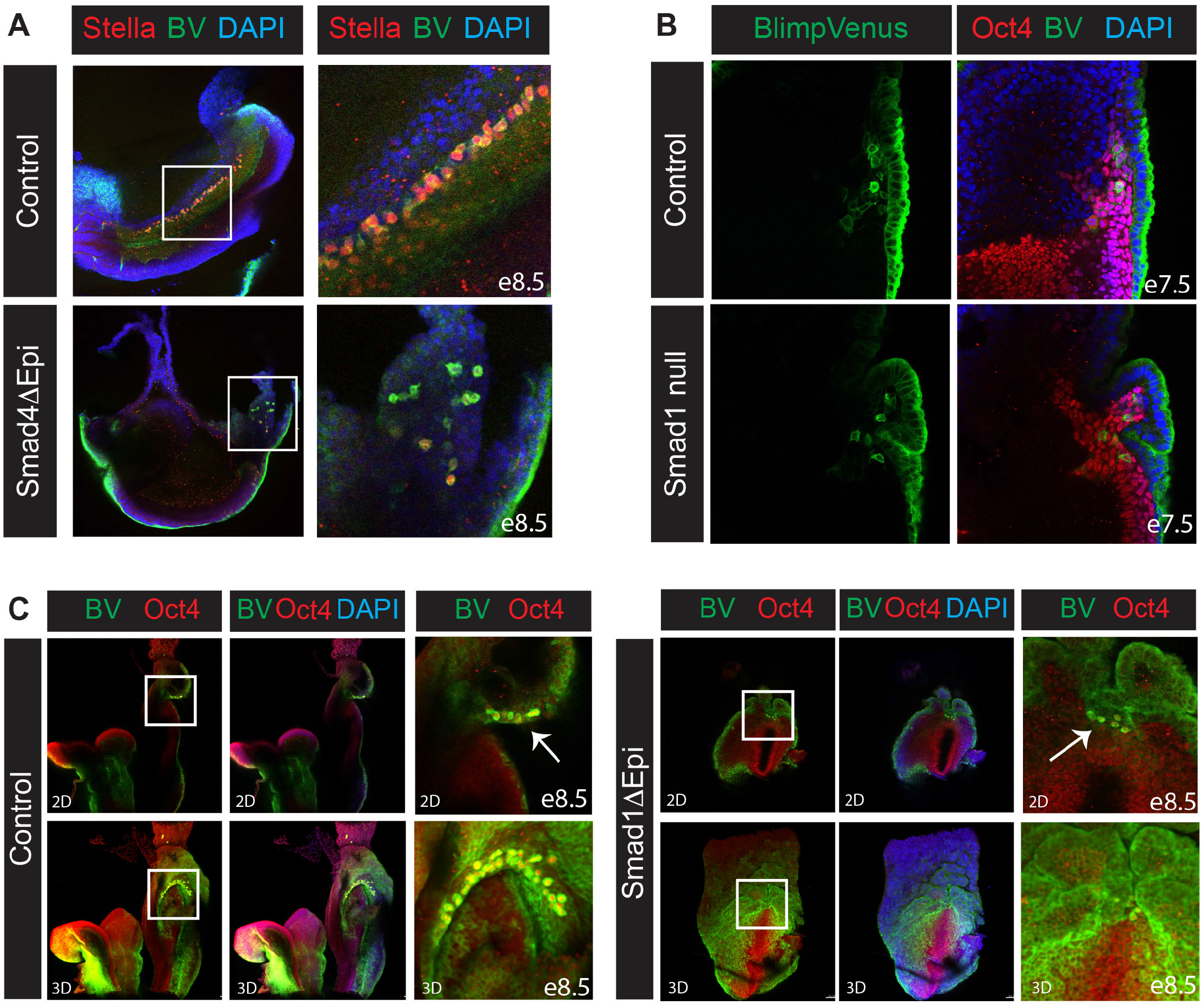
Smad1 is required for PGC specification whereas Smad4 controls PGC maintenance and migration. (A) Stella staining at e8.5 shows PGCs migrating along the hindgut endoderm in BV^+^ control but not Smad4ΔEpi embryos. (B) Oct4 staining in Smad1 null and control e7.5 BV^+^ embryos. (C) Single optical sections (2D) and z-stack projections (3D) showing Oct4 staining in the posterior side of Smad1ΔEpi and control BV^+^ embryos. Arrows indicate Oct4 and BV co-expressing cells. All IF images are counterstained with DAPI.

Smad1^−/−^ embryos form only a rudimentary allantoic bud and develop a distinctive out-pocketing of the proximal posterior VE^15^. At e7.5, only a few BV/Oct4^+^ pre-PGCs are detectable underneath the ruffled VE (Fig. 3B) and mature Stella positive PGCs fail to emerge. To further investigate Smad1 functional requirements in the VE, we used the Ttr-Cre deleter strain. Smad1∆VE embryos display extensive ruffling of the VE (Supplemental Fig. 3D). However, Blimp1^+^ pre-PGCs are occasionally observed in the early epiblast (Supplemental Fig. 3E). An epiblast specific Smad1 deletion likewise results in VE ruffling and the appearance of rare BV/Oct4 double positive cells underneath the overgrown VE (Fig. 3C). However, as for *Smad1* null embryos these BV/Oct4^+^ cells fail to form specified PGCs. Taken together these experiments demonstrate that Smad1 functions in the epiblast during PGC specification, whereas in contrast Smad4 promotes PGC maintenance and/or migration.

### Nodal and Bmp signalling requirements during PGCLC formation *in vitro*

Recent reports have described embryonic stem (ES) cell culture protocols for *in vitro* differentiation of PGC-like cells (PGCLCs), that, as judged by gene expression patterns and global epigenetic remodelling profiles, closely resemble bona fide PGCs^7,8^. To further investigate Nodal and Bmp signalling requirements we generated Smad2^−/−^, Eomes^−/−^, Smad1^−/−^, Smad4^−/−^ and Wnt3^−/−^ ES cell lines carrying the BV-transgene. Control and mutant BV^+^ ES cells, grown under naïve conditions, were induced to form epiblast-like cells (EpiLCs), and subsequently aggregated under appropriate conditions to yield PGCLCs (Fig. 4A). As expected in wild type cultures at day 2 and day 4 of PGCLC differentiation BV/Stella, BV/Oct4, as well as BV/Ap2γ co-expressing cells were detectable by immunofluorescence (Fig. 4B, Supplemental Fig. 4A & 4B). As a negative control, consistent with previous studies^41^, Wnt3^−/−^ cultures contained only a few double positive PGCLCs (Fig. 4B, Supplemental Fig. 4A & 4B). Our Smad2^−/−^, Eomes^−/−^, Smad1^−/−^ and Smad4^−/−^ ES cell lines yielded BV/Stella, BV/Oct4 and BV/Ap2γ double positive PGCLCs (Fig. 4B, Supplemental Fig. 4A & 4B). However, as judged by immunofluorescence and RT-qPCR analysis, the efficiency of PGCLC induction was compromised as compared to wild type control cultures. Thus, Smad2^−/−^, Eomes^−/−^, Smad1^−/−^ and Smad4^−/−^ cell aggregates express lower levels of *Prdm1* (Blimp1) and *Tfap2c* (Ap2γ), and with the exception of Smad2^−/−^ and Eomes^−/−^ cultures, also show increased levels of the somatic marker *Hoxb1* (Fig. 4D). These results demonstrate that outside the context of the early embryo, in cultures containing high levels of exogenously added growth factors, that Eomes^−/−^ and Smad1^−/−^ ES cells can differentiate into PGCLCs.

**Figure 4:**
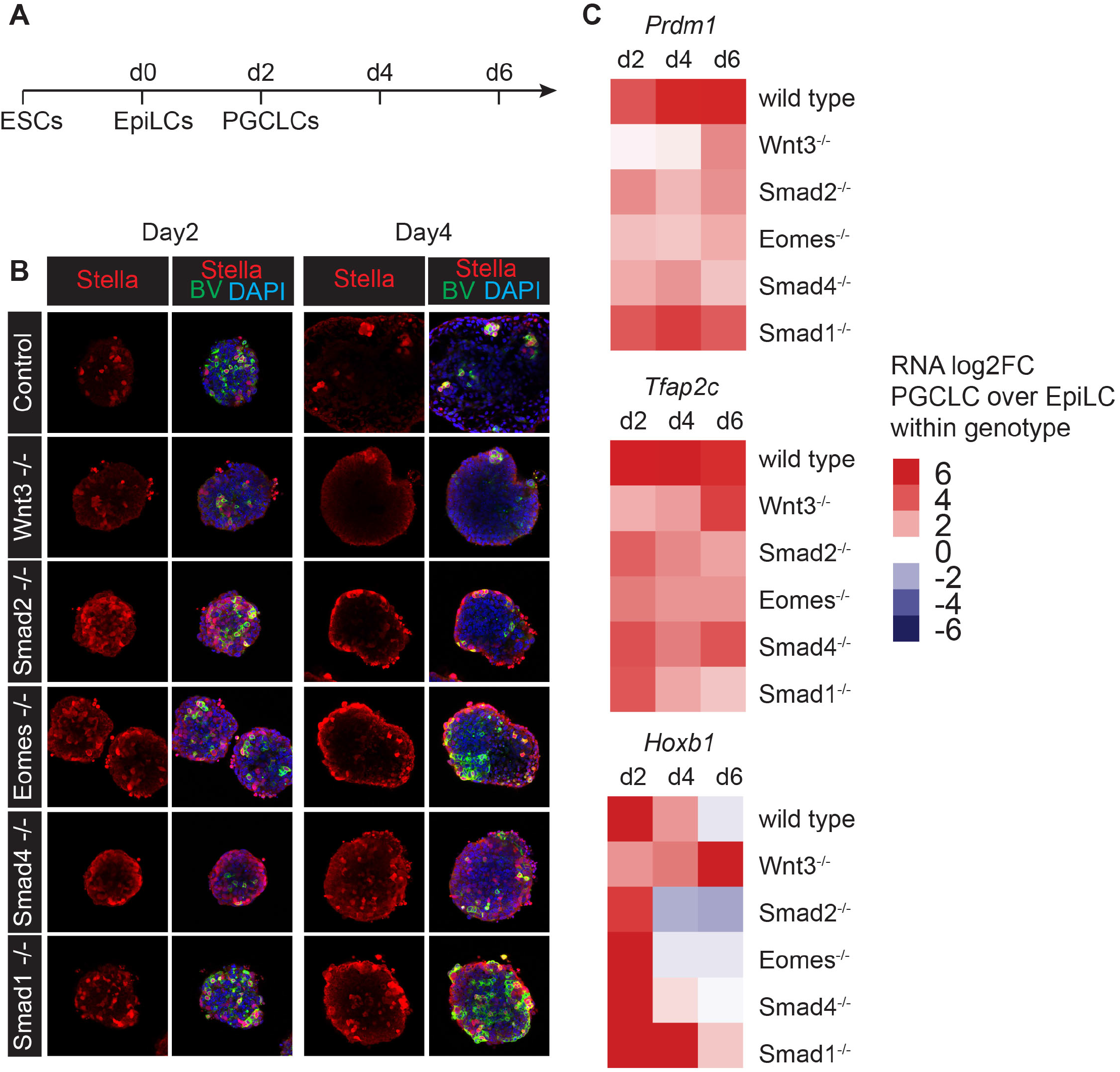
Abilities of mutant ES cells lacking Nodal/Bmp/Smad pathway components to undergo PGCLC differentiation in vitro. (A) Time-line and culture protocol used for PGCLC differentiation. (B) Stella IF staining of day 2 and day 4 BV^+^ PGCLC aggregates derived from the indicated genotypes, counterstained with DAPI (C) Heatmap showing the log2 fold changes (log2FC) of *Prdm1, Tfap2c* and *Hoxb1* expression at day 2, 4 and 6 of PGCLC differentiation in control and mutant cells of the indicated genotypes, relative to EpiLCs.

### Decreased responsiveness to Bmp signalling accompanies PGC specification

At e6.5 p-Smad159 staining is normally restricted to the extra-embryonic VE, the anterior VE and the PPE whereas the posterior embryonic VE lacks p-Smad159 (Fig. 5A). In contrast in e5.5 Smad2∆VE embryos, p-Smad159 is detectable throughout the entire VE (Fig. 1A), but slightly later at e6.5 VE expression is downregulated (Fig. 5A). The mesodermal cell population surrounding BV^+^ cells display strong p-Smad159 immunoreactivity at e6.5. However, the BV^+^ cells themselves are devoid of p-Smad159 staining (Fig. 5A). Likewise, the BV high expressing cells in control e7.5 embryos, as well as the expanded numbers of BV^+^ cells present in e7.5 Smad2∆VE embryos, lack p-Smad159 reactivity (Fig. 5B). These results demonstrate that down-regulated p-Smad159 levels distinguishes specified PGCs from their somatic neighbours and therefore specified PGCs do not respond to high levels of Bmp signalling. Similarly, Eomes∆Epi embryos display a high proportion of BV^+^ epiblast cells lacking p-Smad159 activity (Supplemental Fig. 5A). Decreased p-Smad159 staining was also observed within the BV^+^ cells generated during in vitro PGCLC differentiation (Fig. 5C). Thus, decreased responsiveness to Bmp signalling is a characteristic feature of PGC cell populations, both in *in vitro* and *in vivo* settings.

**Figure 5:**
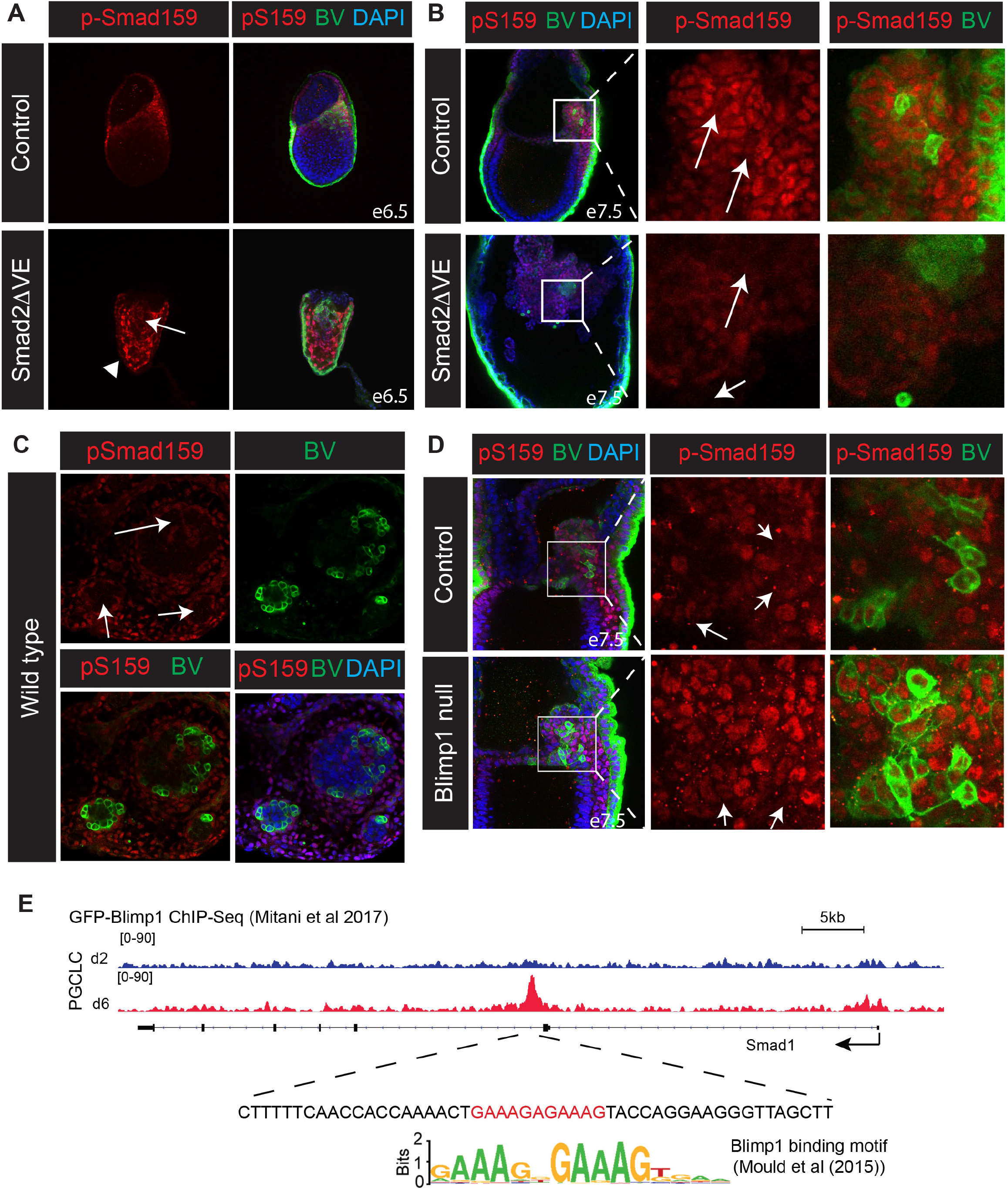
PGCs down-regulate p-Smad159 to become refractory to the high Bmp signalling environment within the proximal posterior epiblast. (A) p-Smad159 staining of BV^+^ cell populations. Arrowhead confirms e6.5 Smad2ΔVE embryos lack p-Smad159 in the VE, while arrow indicates weak activity in BV-expressing epiblast cells. (B). p-Smad159 staining in control and Smad2ΔVE BV^+^ embryos at e7.5. Arrows indicate loss of p-Smad159 staining in BV-high expressing cells. (C) p-Smad159 staining in BV^+^ wild type d6 PGCLC aggregates. Arrows indicate BV^+^ cell populations of lacking p-Smad159 reactivity. (D). p-Smad159 staining in control and Blimp1^−/−^ BV^+^ embryos at e7.5. Arrows indicate p-Smad159 reactivity in BV-high expressing cells in expanded panels. All IF images are counterstained with DAPI. (E) Genome browser track view of GFP-ChIP wiggle plots from GFP-Blimp1 expressing PGCLC cultures at day (d)2 (blue) and d6 (red) showing enrichment of GFP-Blimp1 density at a *Smad1* intronic region. The sequence below the GFP-Blimp1 ChIP site contains a Blimp1 binding motif indicated in red. The consensus Blimp1 binding motif previously identified by genome wide ChIP-seq data sets is also indicated^65^.

To directly assess whether Blimp1 expression influences Smad159 phosphorylation levels, we generated Blimp1^−/−^ embryos carrying the BV-transgene. BV^+^ epiblast cells were visible at the base of the allantoic bud at e7.5 in Blimp1^−/−^ embryos (Fig. 5D), strengthening the notion that Blimp1 functional activity is not required at early stages during pre-PGC formation. Interestingly we found that the BV high cell population partially retains p-Smad159 reactivity (Fig. 5D). To test whether loss of p-Smad159 within the maturing PGCs potentially reflects reduced levels of transcription, next we examined RNA-seq data sets^42,43^. Reduced *Smad1* and *Smad5* expression levels were reported in e7.5 PGCs as compared to the surrounding mesodermal cell population (Supplemental Fig.5B). Moreover RNA-seq analysis of wild type PGCs confirms that *Smad1* expression is down-regulated during PGC development (Supplemental Fig. 5C), whereas in contrast BV^+^ Blimp1^−/−^ PGCs fail to down-regulate *Smad1* expression (Supplemental Fig. 5C). Consistent with this recent Blimp1 ChIP-Seq experiments analysing d6 PGCLCs^43^ allowed us to identify a Blimp1 peak downstream of the *Smad1* first coding exon containing the canonical Blimp1 binding motif (Fig. 5E). Collectively these observations strongly suggest that Blimp1 may directly repress *Smad1* transcription during early PGC development. Conditional deletion of Blimp1 in mature PGCs, using the Stella-Cre deleter strain^43^, has no effect on *Smad1* transcriptional levels (Supplemental Fig. 5C). Thus, Blimp1 downregulates *Smad1* expression at early stages of PGC development ensuring that the emerging PCGs are refractory to the high Bmp signalling environment in the posterior epiblast that normally promotes development of the ExM.

## Discussion

The transcriptional regulators and epigenetic machinery responsible for guiding allocation of the PGC lineage have been intensively investigated over the past decade^1,5^. A small population of pre-PGCs (~5) initially present in the proximal epiblast in pre-streak e6.25 embryos are marked by co-expression of the zinc finger transcriptional repressor Blimp1^3,44^, together with the pluripotency factors Oct4 and Nanog. Slightly later at e6.75, *Prdm14* expression is induced in developing PGCs^45^ and subsequently, coincident with formation of the posterior ExM, PGCs activate expression of *Dppa3* (Stella), *Tfap2c* (Ap2γ) and *Sox2*. The cluster of specified PGCs (~40) is detectable at e7.5 at the base of the allantois within the ExM. These committed PGCs then migrate into the endodermal layer of the forming hindgut pocket over the following 24 hours.

Previously we characterised dose-dependent Nodal/Smad functional activities required for A-P and left-right axis patterning in the early mouse embryo^10^. Functional contributions during gastrulation and within the PS derivatives have been extensively characterised^10^. Here we dissect the Nodal/Bmp/Smad signalling cues in the embryonic and extra-embryonic tissues governing the PCG developmental programme. The present experiments demonstrate for the first time that Nodal/Smad2 activities regulate allocation of the pre-PGC population. Thus Nodal null embryos prematurely initiate pre-PGC formation at e5.5. However, these BV^+^ epiblast cells are ectopically positioned and lack Nanog expression. Nanog has previously been shown as an important regulator of PGC development *in vivo*^46,47^, and can induce PGCLC differentiation in EpiLCs *in vitro*^48^. Nanog directly upregulates *Prdm1* (Blimp1) and *Prdm14* expression in PGCLCs^48^. However, Nanog over-expression in EpiLCs activates *Prdm14* expression before *Prdm1*^48^, whereas normally *Prdm1* expression in pre-PGCs precedes that of *Prdm14*^45,48^. These results suggest that Nanog re-expression downstream of Nodal is non-essential for induction of *Prdm1* expression *per se*. Rather Nanog may function primarily to activate *Prdm14*. Here we found that high levels of Nodal signalling in the proximal epiblast, prior to the onset of streak formation, are required to fully reactivate and maintain high levels of Nanog expression.

Nodal is transiently expressed in the proximal epiblast cell population but expression decreases as the streak elongates^49^. Highest levels retained in the anterior streak function to induce DE and midline fates^10^. Mutant embryos lacking the Nodal downstream effector Smad2 fail to induce the Nodal antagonists *Cer1* and *Lefty1* in the VE and consequently display ectopic Nodal signalling throughout the entire epiblast^12,25^. The present experiments demonstrate that conditional Smad2∆VE mutants phenocopy Smad2 null embryos. As judged by downregulated expression of Otx2 and Sox2, together with upregulated Brachyury expression, the epiblast appears to prematurely adopt a proximal posterior character. An inverse relationship between p-Smad159 and p-Smad23 signalling in the VE has previously been reported^9^. Here we found that e5.5 Smad2∆VE embryos display ectopic p-Smad159 activation throughout the VE. Interestingly expanded Nodal activity in the epiblast results in decreased expression of the Bmp antagonist *Gdf3* and likely accounts for the increased levels of p-Smad159 in the VE. Imbalanced Nodal/Bmp signalling leads to the premature appearance and markedly increased numbers of pre-PGCs at e6.5. These results demonstrate Nodal/Smad2 signals localised to the proximal epiblast govern competency of the proximal epiblast to acquire PGC characteristics and promote the establishment of the PGC niche.

Recent work suggests that occupancy at a putative T-box site, 10kb upstream of the *Prdm1* transcription start site, by the T-box transcription factor family member Brachyury activates Blimp1 expression in the pre-PGC population^41^. Consistent with this, Brachyury mutant embryos activate but fail to maintain BV^+^ pre-PGCs^41^. However, forced expression of either Brachyury or Eomes in EpiLCs leads to up-regulated *Prdm1* expression^41^. Eomes and Brachyury could potentially function collaboratively to regulate *Prdm1* expression. However, the present experiments demonstrate that Eomes is dispensable for initial Blimp1 induction in the epiblast. Moreover, the BV^+^ cells formed in the absence of Eomes mostly lack Brachyury expression. These observations strongly suggest that Brachyury and Eomes are dispensable for induction of Blimp1.

Eomes in the early PS directly activates *Mesp1* expression necessary to allow nascent mesoderm to down-regulate E-Cadherin and undergo EMT^36,50^. Here we demonstrate that Eomes, acting immediately downstream of Nodal within the epiblast^37^, plays an essential role during the early stages of PGC formation. In the absence of Eomes, we observe an expanded population of BV^+^ cells in the proximal epiblast. However, these pre-PGC-like cells lack Nanog expression and fail to activate the germ cell programme. Interestingly Eomes mutant BV^+^ cells, in close proximity to Bmp4 signals from the overlying ExE, lack p-Smad159 activity and are refractory to Bmp signalling. The absence of Eomes in the proximal epiblast disturbs formation of the PGC niche within the posterior ExM population that normally promotes PGC specification and survival^51,52^. Previous work shows that Eomes null primitive streak cells can undergo EMT and generate mesodermal cell populations *in vitro*^36^. Here we demonstrate that Eomes null ES cells can differentiate into BV/Oct4/Ap2γ/Stella positive PGCLCs, albeit at a lower efficiency as compared to wild-type controls, most likely due to their ability to generate a supporting mesodermal population. Thus, Eomes is not intrinsically required for Blimp1 induction and PGC formation per se. Rather, acting downstream of Nodal, Eomes is essential to promote formation of the posterior signalling niche necessary for maturation of the PGC lineage.

How PGCs maintain their unique cell-type identity within the predominant mesodermal signalling environment at the base of the allantois remains mysterious. Here we describe a novel cellular mechanism that allows PGCs to insulate themselves and become refractory to Bmp signals. Thus, we found at e7.5 that BV^+^ cells display reduced Smad159 phosphorylation levels. Moreover within d6 PGCLC aggregates p-Smad159 activity and BV expression are mutually exclusive. Downregulated expression of both *Smad1* and *Smad5* was previously observed in PGCs compared to the surrounding mesoderm^42^. Blimp1 may directly repress *Smad1* transcription via occupancy at a site within the Smad1 locus. Consistent with this idea recent RNA-seq data sets^43^ reveal that *Smad1* expression in Blimp1-deficient pre-PGCs is not down-regulated.

The cluster of PGCs at the base of the allantois, that themselves lack Bmp4 expression, are surrounded by ExM expressing high levels of Bmp4^17^. While PGCs themselves become refractory to the Bmp signalling environment, their survival and further development is thought to be critically dependent on the ability of the adjacent ExM to provide extrinsic trophic growth factors^52^. Recent *in vitro* experiments similarly suggest that the role of Smad1 signalling during PGCLC differentiation may be to generate somatic cells that function to promote PGCLC survival^53^. Likewise, we observe here in PGCLC aggregates localised pockets of BV^+^ cells within p-Smad159 active regions. Thus, a continuum of Bmp signalling in the early post-implantation embryo is required to induce the formation of pre-PGCS in the early epiblast and subsequently at later stages establishes the posterior signalling niche necessary for PGC specification.

Imaging studies demonstrate that PGCs actively migrate as individual cells from the base of the allantois towards the overlying endoderm^54^. However, the molecular pathways guiding this initial directional migration remain ill-defined. Besides intrinsic regulators of cell motility the posterior endoderm also plays a critical role guiding PGC migration. For example, in Sox17 mutant embryos that display defective endoderm formation, PGCs are formed, mature and initiate migration towards the endoderm but lack the ability to appropriately integrate to the endoderm^55^. Similarly we found here in Smad2∆VE and Smad4∆Epi embryos that PGCs are appropriately specified but fail to migrate towards the endoderm. PGCs normally extend long filopodia-like structures at e7.5^54^. In contrast in Smad2∆VE embryos endoderm formation is compromised, filopodia-like structures are never observed and PGCs remain as clusters of epithelial-like cells. Likewise, Smad4∆Epi embryos fail to form DE^39^. Migration defects could potentially reflect the intrinsic loss of Smad4-dependent PGC functional activities, or alternatively, defective endoderm formation potentially results in the failure to produce chemo-attractants. The extracellular matrix (ECM) also plays a crucial role in guiding PGC migration^56,57^. We previously reported that Smad4 controls ECM deposition in early developmental stages^58^. Further studies on the trophic signals emitted from the posterior endoderm and the potential role of Smad4-dependent signalling in ECM composition will be required to further define Bmp/Nodal Smad requirements during PGC migration.

An antagonistic relationship between Nodal and Bmp pathways has been described in a variety of developmental contexts including within the VE for establishing initial proximal-distal polarity, patterning of the PS, morphogenesis of the amnion and correct establishment of the left-right body plan^9,10,13,59,60^. The present experiments further demonstrate that these regulatory cues govern cell fate decisions in the posterior epiblast causing a discrete subpopulation to adopt a germ cell versus somatic cell fate. Importantly, we found that Blimp1 induction within the PGCs is associated with down-regulated pSmad159 levels and thus provide a cell intrinsic regulatory mechanism that allows this subpopulation to become non-responsive to local Bmp signalling cues. Future experiments will be needed to further define dynamic cellular events controlling this developmental switch.

## Methods

### Animal care

*Smad2*^−/− 25^, *Smad2^CA^* ^61^, *Nodal*^−/−^ *^24^*, *Eomes*^−/−^, *Eomes^CA^* ^36^, *Smad4*^−/−^, *Smad4^CA^* ^39^, *Prdm1*^−/−^ *^4^*, *Prdm1^CA^* ^62^, *Smad1*^−/−^, *Smad1^CA^* ^15^, *Prdm1^Cre-Lacz^* ^40^, *Ttr-Cre* ^26^, *Sox2-Cre*^33^, *Prdm1-mVenus* ^32^ alleles were genotyped as described. Supplemental Table S2 indicates inter-crosses produced for this study and acronyms used for embryos. All animal experiments were performed in accordance with Home Office (UK) regulations and approved by the University of Oxford Local Ethical Committee.

### Generation of knockout ES cell lines and ES cell culture

Blastocysts for ESC derivation were obtained from inter-crosses of *Wnt3^+/−^* ^23^, *Smad1^+/− 15^*, *Smad2^+/−^* ^25^, *Smad4^+/^−* ^39^ and *Eomes^+/−^* ^36^ mice harbouring the *Prdm1.Venus* BAC transgene ^32^. All ESC lines used were grown in feeder-free conditions on 0.1% gelatin-coated dishes at 6% CO2 at 37°C. ESCs were cultured in serum-free media containing N2B27 (Takara, Y40002) supplemented with 1μM PD0325091 and 3μM CHIR99021 and 1000 U/ml LIF.

### PGCLC cultures

EpiLCs and PGCLCs were induced as previously described^63^. In brief, 7 × 10^5^ ESCs were washed and resuspended in N2B27 medium (Takara, Cat#Y40002) supplemented with 12ng/ml bFgf (Invitrogen, 13256-029), 20ng/ml Activin A and 1% KSR (Gibco, 10828, Lot:1508151) and grown on fibronectin-coated (10μg/cm^2^) (Millipore, FC010) 6cm dishes for 48h to form EpiLCs. 2000 EpiLCs were then washed and plated into ultra-low attachment U-bottom shaped 96-well plates in serum-free medium (GK15; GMEM (Invitrogen) with 15% KSR, 0.1 mM NEAA, 1 mM sodium pyruvate, 0.1mM 2-mercaptoethanol, 100 U/ml penicillin, 0.1 mg/ml streptomycin, and 2 mM L-glutamine) in the presence of the cytokines BMP4 (500 ng/ml; R&D Systems, 314-BP), LIF (1000 u/ml; Millipore, ESG1107), SCF (100 ng/ml; R&D Systems, 455-MC), BMP8b (500 ng/ml; R&D Systems, 1073-BP), and EGF (50 ng/ml; R&D Systems, 2028-EG).

### Immunofluorescence

d2 and d4 PGCLCs as well as E5.5-E8.5 mouse embryos were harvested, washed in PBS and fixed in 1% paraformaldehyde (PFA) o/n at 4°C. After three washes in PBS containing 0.1% Triton X-100 (PBS-Tx), samples were permeabilised in PBS containing 0.5% Triton X-100 followed by three washes in PBS-Tx, blocked in 5% donkey serum and 0.2% BSA in PBS-Tx for 1h at RT and incubated overnight with primary antibodies in blocking solution at 4°C. Following four washes in PBS-Tx, samples were incubated with fluorophore-conjugated secondary antibodies in blocking solution (2h, RT) followed by three-five washes in PBS-Tx, one wash in PBS-Tx containing 2μg/ml DAPI and three washes in PBS-Tx prior to mounting in Vectashield with DAPI (H-1200). Samples were imaged the following day on an Olympus Fluoview FV1000 confocal microscope and image data was processed using ImageJ and Bitplane Imaris software. Antibodies are listed in the Supplementary Table S3.

### RT-qPCR

250ng RNA was reverse transcribed to cDNA using Superscript III First Strand Synthesis System (Life Technologies, Cat#18080-051) and diluted to 100μl final volume in H2O (2.5ng/ul). 2μl (5ng) cDNA were used per RT-qPCR reaction in duplicate using SYBR-green kit (Qiagen, Cat#204143). Relative gene expression was normalised to *Gapdh* expression and calculated as 2^ΔΔ^Ct. Average expression levels were calculated for two technical replicates from two independent cell lines per genotype. RT-qPCR primer sequences are listed in Table S4.

### *In situ* hybridisation and immunohistochemistry

Whole-mount *in situ* hybridisation analysis was performed as before^50^, using antisense riboprobes for *Bmp2*^64^, *Bmp4*^13^ and *Gdf3*^28^. For Blimp1 immunohistochemistry, E7.5 decidua were fixed overnight in 4% PFA, dehydrated in ethanol, embedded in paraffin wax and sectioned (6 μm). Dewaxed sections were subjected to antigen retrieval by boiling for 1h in Tris/EDTA (pH 9.0) and permeabilized for 10 min in 0.1% Triton X-100 in TBS. Sections were subsequently blocked with 10% normal goat serum in TBS. Sections were incubated in rat monoclonal anti-Blimp1 (1:500 dilution, sc-130917, Santa Cruz Biotechnology) in block overnight at 4°C and signal-amplified with rabbit anti-rat secondary antibody (AI-4001, Vector Laboratories) for 45min at RT followed by peroxidase blocking for 20 minutes at RT and development with Envison System-HRP for rabbit antibodies (K4011, DAKO) and Vector Red substrate (SK-4805, Vector Laboratories). Sections were lightly counter-stained with haematoxylin, coverslipped and imaged. Haematoxylin and eosin staining was performed as previously described^36^.

## Acknowledgements

We would like to thank Mitinouri Saitou for generously providing the Blimp1-mVenus BAC transgenic mouse strain. Confocal microscopy was carried out in the Micron Advanced Bioimaging Unit (funded by Wellcome Trust Strategic Award 107457). This work was funded by the Wellcome Trust (099840/Z/12/A to A.D.S. and 102811/Z/13/2 to E.J.R.). E.J.R. is a Wellcome Trust Principal Research Fellow.

## Author Information

All the authors designed the experiments, A.D.S. and I.C. performed the experiments. All of the authors contributed to writing the paper.

## Supplemental Figure Legends

**Supplemental Figure 1:**
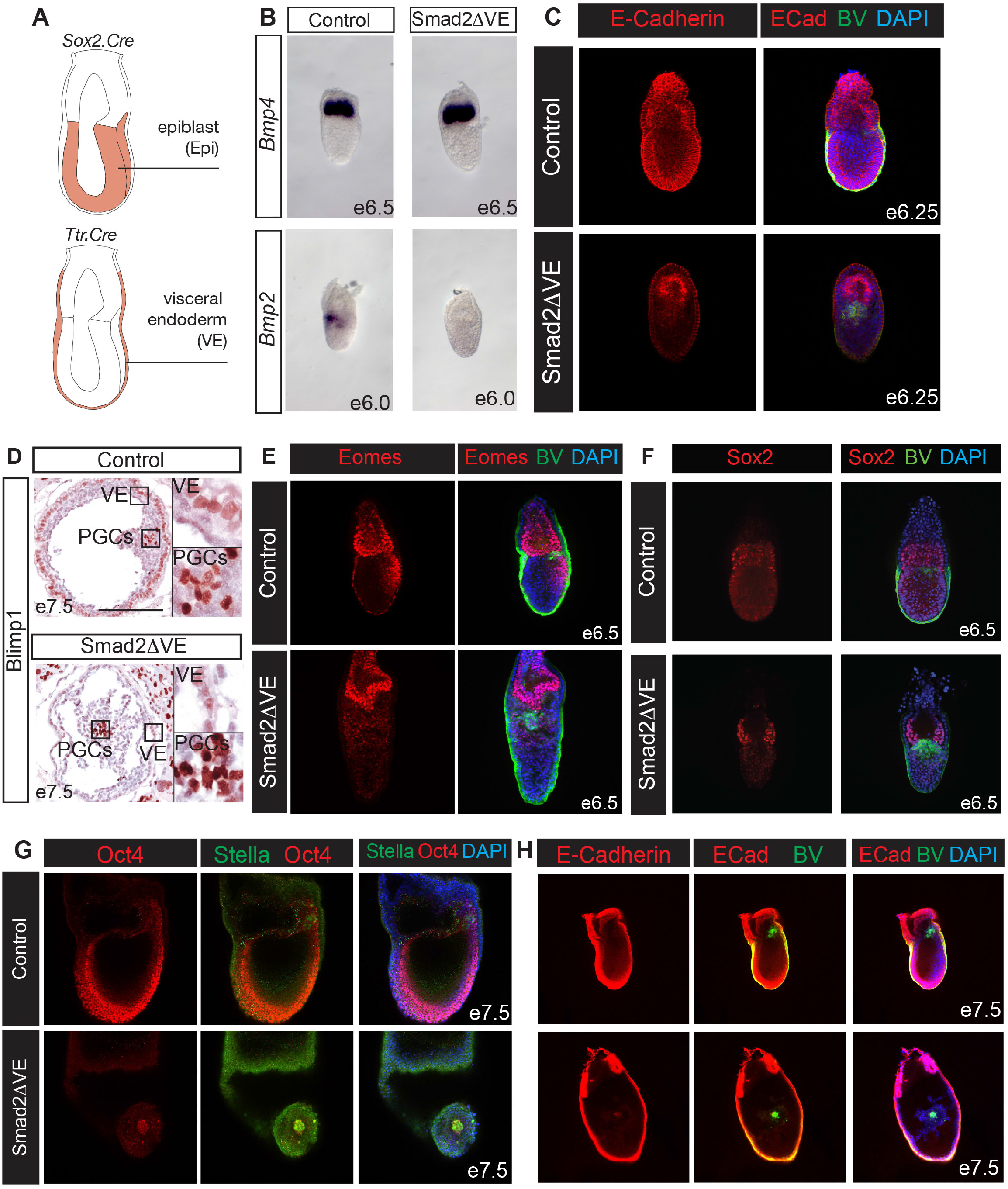
Conditional Smad2 inactivation in the visceral endoderm results in expansion of the PGC niche. (A) Tissue specific activity of the Sox2 and Ttr Cre deleter strains. (B) Whole-mount *in situ* hybridisation analysis of *Bmp4* and *Bmp2* expression in e6.5 and e6.0 control and Smad2ΔVE embryos. (C) E-Cadherin (ECad) staining of e6.25 control and Smad2ΔVE BV^+^ embryos. (D) Immunohistochemistry (IHC) staining of sections of e7.5 control and Smad2ΔVE embryos using a Blimp1 antibody. Boxed regions are expanded on the right. Scale bar = 200µm. (E) Eomes expression in control and Smad2ΔVE BV^+^ e6.5 embryos. (F) Sox2 expression in BV^+^ e6.5 embryos. (G) Stella and Oct4 co-localization in e7.5 control and Smad2ΔVE embryos. (H) ECadherin staining of e7.5 control and Smad2ΔVE embryos. All IF images are counterstained with DAPI.

**Supplemental Figure 2:**
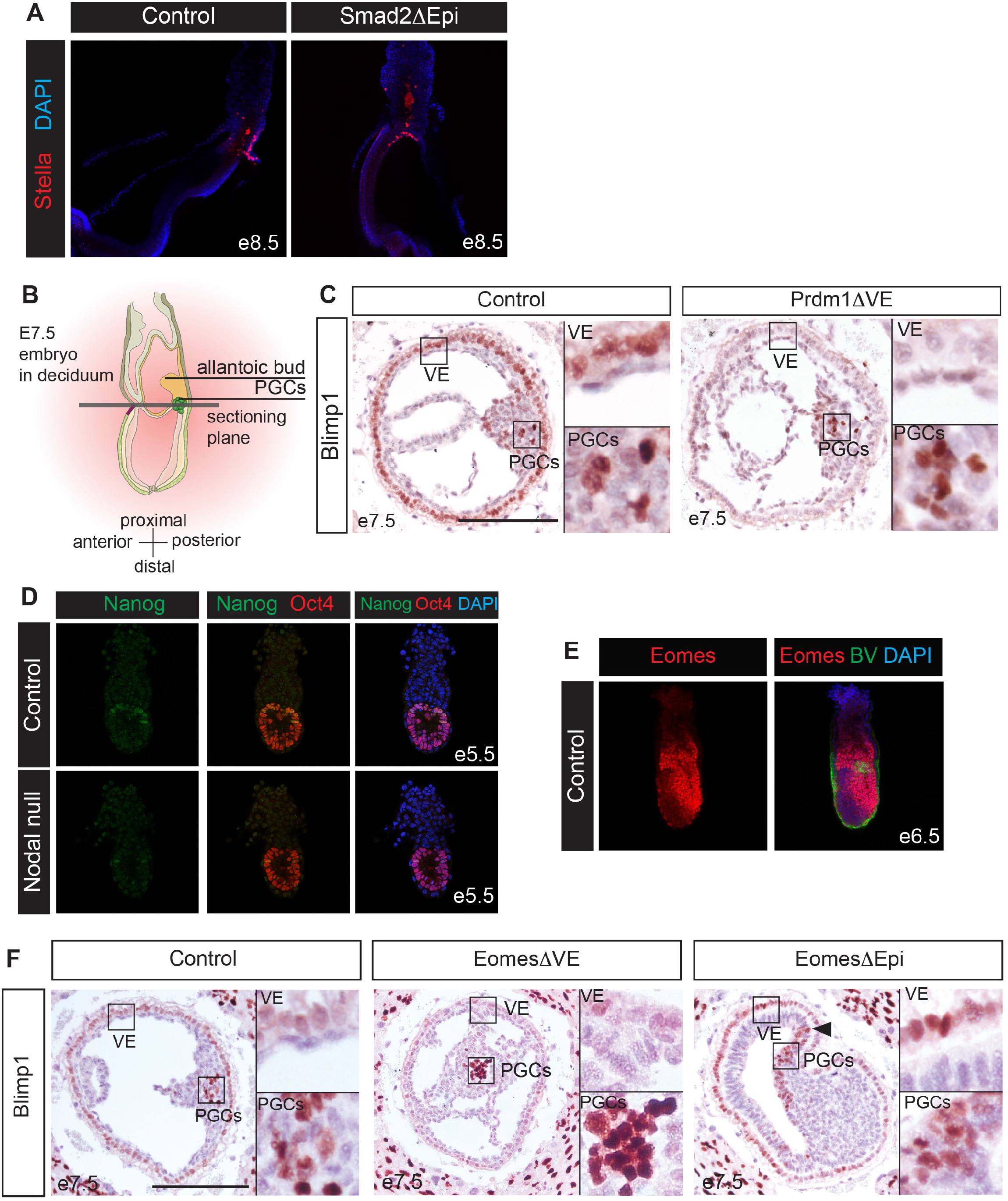
Smad2, Prdm1, Nodal and Eomes functional contributions during PGC development. (A) Stella IF staining in control and Smad2ΔEpi e8.5 embryos. (B) Plane of section used for IHC staining. (C) Blimp1 IHC in control and Prdm1ΔVE sections shows selective deletion of Blimp1 in the VE of Prdm1ΔVE embryos fails to prevent PGC formation. Scale bar = 200µm. (D) Nanog and Oct4 staining in e5.5 control and Nodal null embryos. (E) Eomes staining in mid-late streak stage control BV^+^ embryos. (F) Blimp1 IHC in e7.5 control, EomesΔVE and EomesΔEpi embryos. Boxed areas indicated expanded panels on the right of the image. Scale bar = 200µm.

**Supplemental Figure 3:**
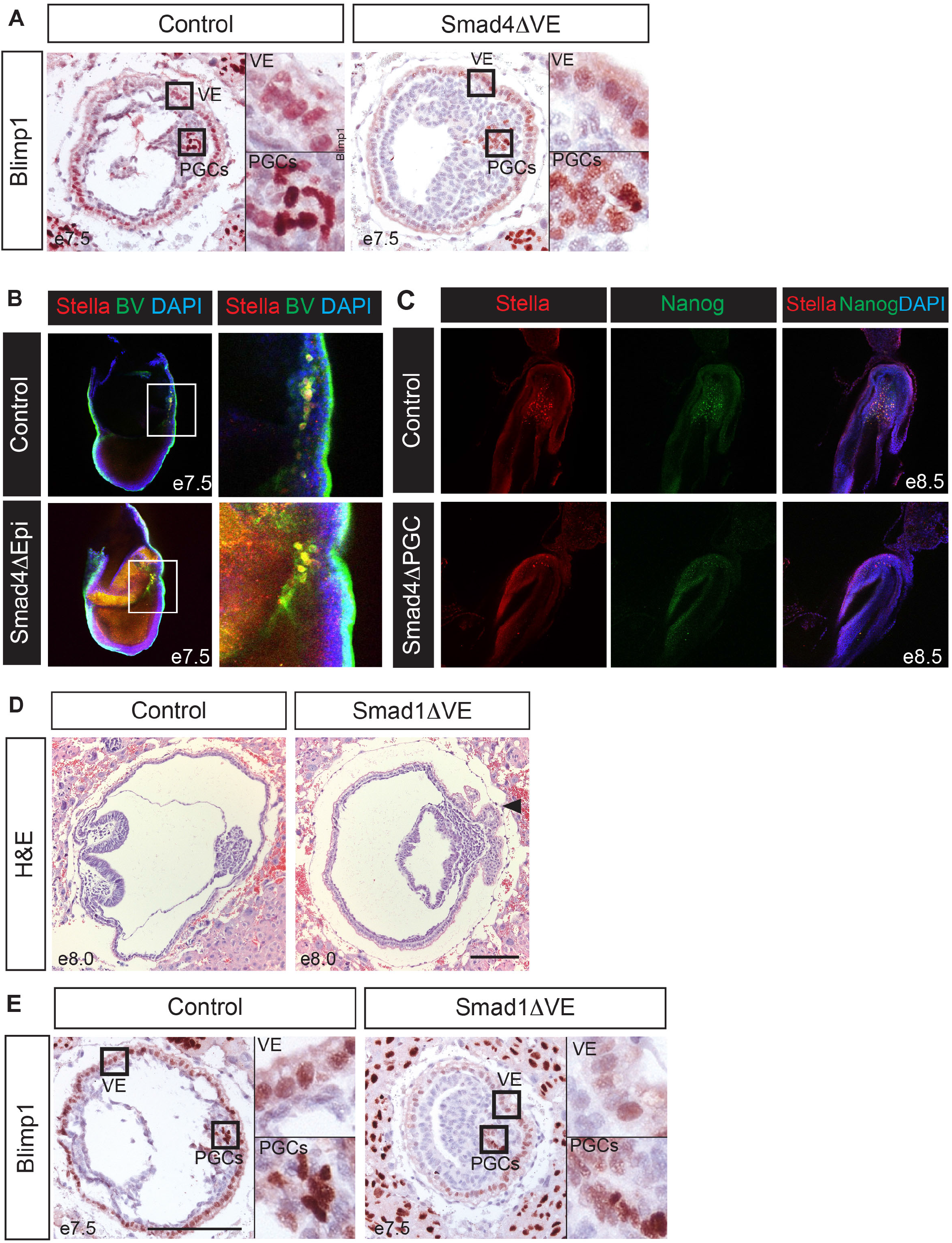
Smad4 and Smad1 requirements for promoting PGC development. (A) Blimp1 IHC staining of e7.5 control and Smad4ΔVE embryo sections. Boxed areas indicated expanded panels on the right of the image. (B) Stella expression in e7.5 control and Smad4ΔEpi BV^+^ embryos. (C) Optical sections of the posterior region of e8.5 control and Smad4ΔPGC embryos co-stained with Stella and Nanog antibodies, counterstained with DAPI. (D) H&E staining of e8.0 control and Smad1ΔVE embryo sections. Arrowhead indicates ruffling of the posterior VE in Smad1ΔVE embryos. (E) Blimp1 IHC in control and Smad1ΔVE e7.5 embryo sections. Boxed areas indicated expanded panels on the right of the image. Scale bar = 200µm.

**Supplemental Figure 4:**
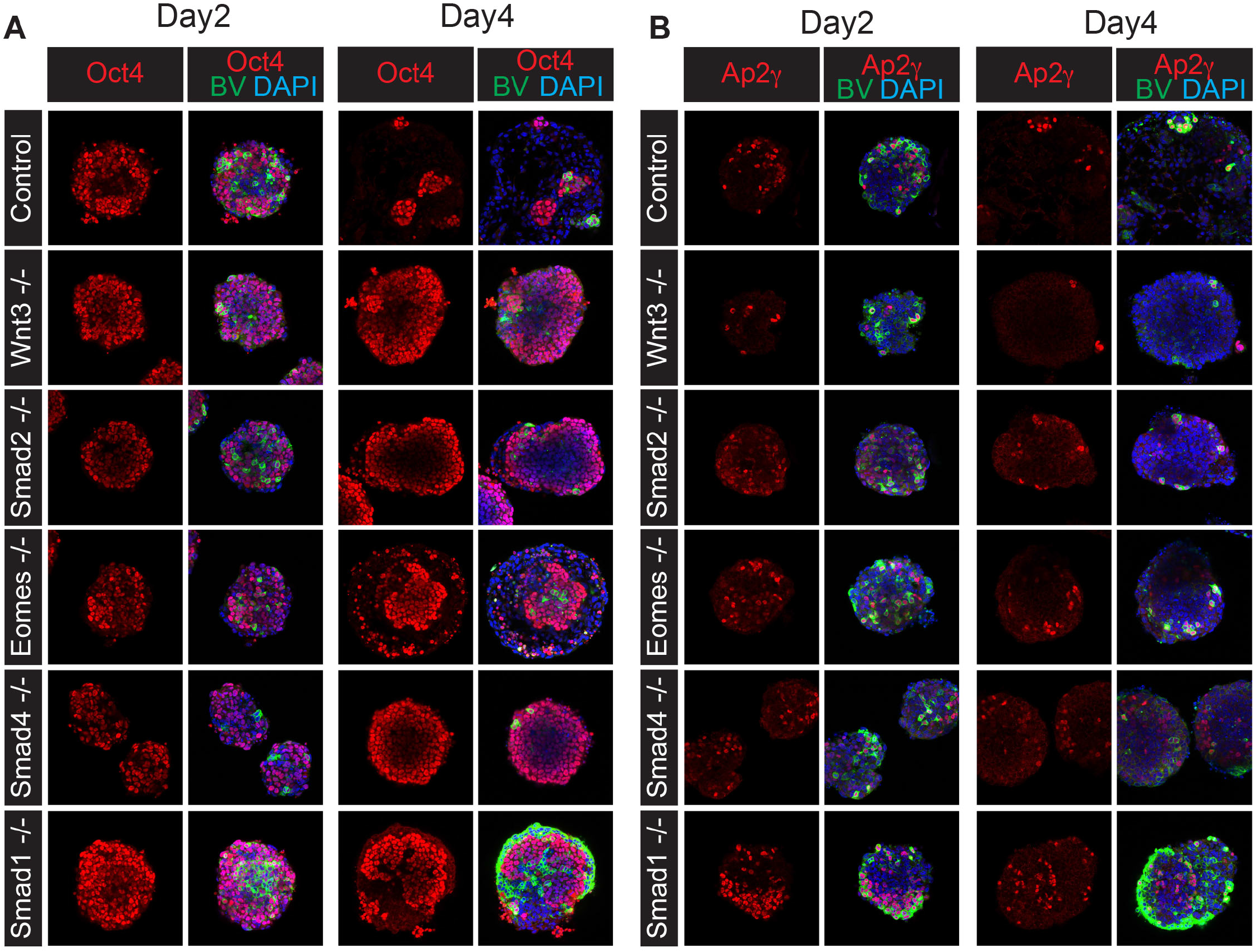
Expression of PGC marker gene expression in wild type compared to Nodal, Bmp and Wnt signalling mutant PGCLC aggregates. (A) Oct4 staining of day 2 and day 4 PGCLC BV^+^ aggregates of indicated genotypes, counterstained with DAPI. (B) Ap2γ staining of day 2 and day 4 PGCLC BV^+^ aggregates of indicated genotypes, counterstained with DAPI.

**Supplemental Figure 5:**
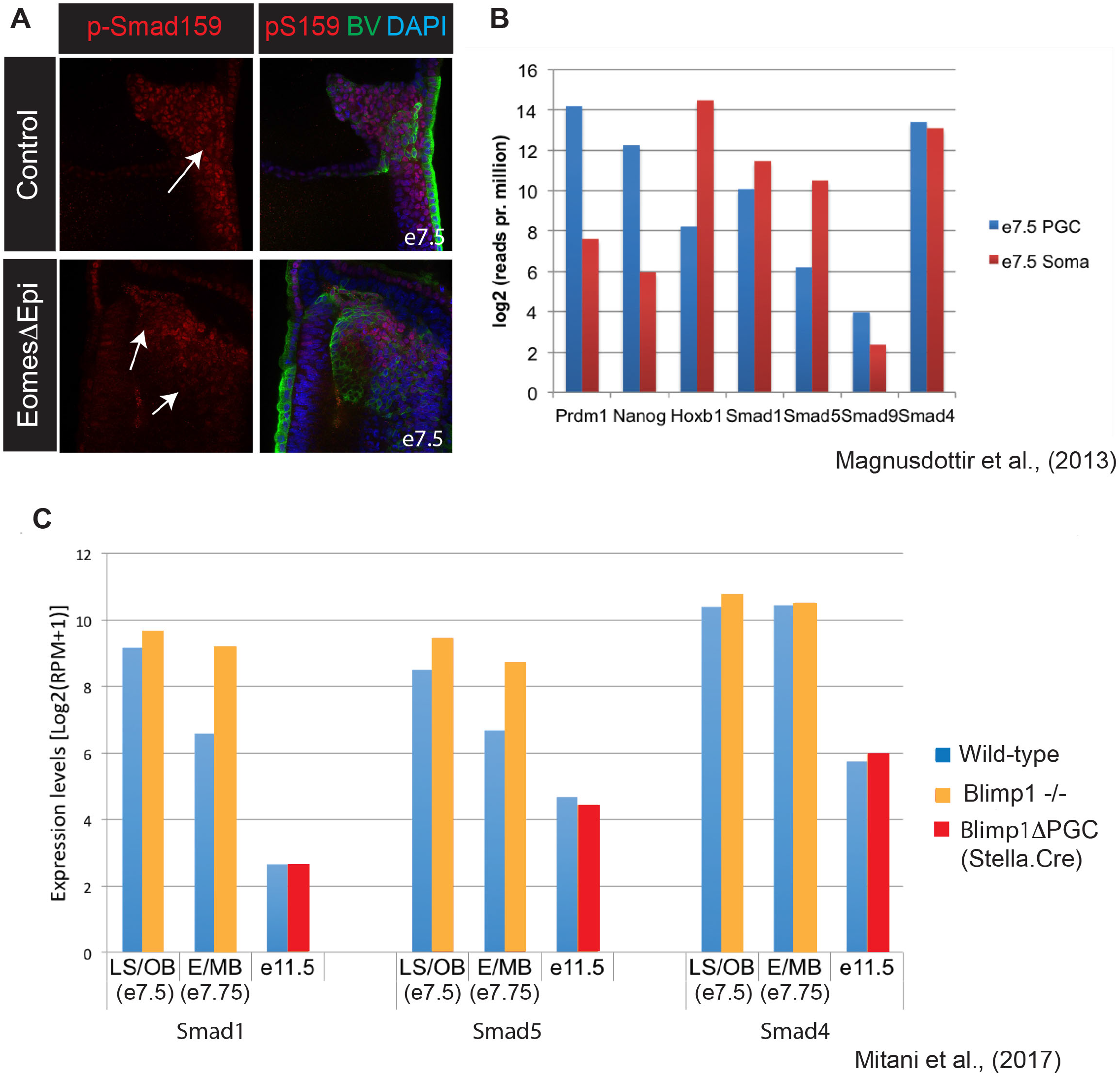
PGCs selectively repress expression of Bmp/Smad signalling components. (A) p-Smad159 staining in e6.5 control and EomesΔEpi BV^+^ embryos. Arrows indicate loss of p-Smad159 signal in the BV^+^ cell population (B) RNA-seq data showing expression levels [log2 (reads per million)] of *Smad1*, *Smad5, Smad9* and *Smad4* in e7.5 PGCs (blue) compared to somatic cells (red)^42^. Expression levels of *Prdm1* and *Nanog* (PGC markers) and Hoxb1 (somatic cell marker) are shown for comparison. (C) RNA-Seq data at indicated time-points showing expression [Log2(reads per million +1)] of *Smad1, Smad5* and *Smad4* in BV^+^ PGC cells isolated from wild type (blue), Blimp1 null (orange) or embryos where Blimp1 has been deleted from specified PGCs using a Stella-Cre deletor strain (Blimp1ΔPGC) (red)^43^.LS/OB, late streak/no bud stage embryo; E/MB, early/mid bud stage embryos.

**Supplemental Table S1.** A. Genotypes of offspring from conditional deletion of *Prdm1* from VE

| *Prdm1*<sup>-/+</sup>; *Ttr-Cre*<sup>+</sup> x *Prdm1*<sup>CA/CA</sup> |  |  |  |  |
| --- | --- | --- | --- | --- |
| Genotype | *Prdm1*<sup>+/<sup>CA</sup></sup> | *Prdm1*<sup>+/<sup>ΔVE</sup></sup> | *Prdm1*<sup>m/<sup>CA</sup></sup> | *Prdm1*<sup>m/<sup>ΔVE</sup></sup> |
| No of animals | 2 | 3 | 3 | 3 |
| Percentage | 18.2 | 27.3 | 27.3 | 27.3 |
| Expected % | 25 | 25 | 25 | 25 |

B. Viability test of *Prdm1*ΔVE intercross
| *Prdm1*<sup>m/<sup>ΔVE</sup></sup> x *Prdm1*<sup>m/<sup>ΔVE</sup></sup> |  |
| --- | --- |
| No of animals | 9 |

**Supplemental Table S2.**

| Embryo name | Genotype | Phenotype<br>Reference | Additional transgene<br>(Ohinata et al. 2008) |
| --- | --- | --- | --- |
| Smad2ΔVE | *Smad2<sup>CA/-</sup> ; Ttr.Cre<sup>+/-</sup>* |  | *Blimp1.Venus* |
| Smad2ΔEpi | *Smad2<sup>CA/-</sup> ; Sox2.Cre<sup>+/-</sup>* | Vincent et al. 2003 |  |
| EomesΔVE | *Eomes<sup>CA/-</sup> ; Ttr.Cre<sup>+/-</sup>* | Nowotschin et al. 2013 |  |
| EomesΔEpi | *Eomes<sup>CA/-</sup> ; Sox2.Cre<sup>+/-</sup>* | Arnold et al. 2008 | *Blimp1.Venus* |
| Smad1ΔVE | *Smad1<sup>CA/-</sup> ; Ttr.Cre<sup>+/-</sup>* |  |  |
| Smad1ΔEpi | *Smad1<sup>CA/-</sup> ; Sox2.Cre<sup>+/-</sup>* |  | *Blimp1.Venus* |
| Smad1 null | *Smad1<sup>-/-</sup>* | Tremblay et al. 2001 | *Blimp1.Venus* |
| Smad4ΔVE | *Smad4<sup>CA/-</sup> ; Ttr.Cre<sup>+/-</sup>* | Li et al. 2010 |  |
| Smad4ΔEpi | *Smad4<sup>CA/-</sup> ; Sox2.Cre<sup>+/-</sup>* | Chu et al. 2004 | *Blimp1.Venus* |
| Smad4ΔPGC | *Smad4<sup>CA/-</sup> ; Blimp1.iCre<sup>+/-</sup>* |  | *Blimp1.Venus* |
| Prdm1ΔVE | *Prdm1<sup>CA/-</sup> ; Ttr.Cre<sup>+/-</sup>* |  |  |
| Prdm1 null | *Prdm1<sup>-/-</sup>* | Vincent et al. 2005 | *Blimp1.Venus* |
| Nodal null | *Nodal<sup>-/-</sup>* | Conlon et al. 1991 | *Blimp1.Venus* |

Sox2.Cre reference: Hayashi et al. 2002. Ttr.Cre reference: Kwon and Hadjantonakis 2009.

**Supplemental Table S3.**

| Antibodies | SOURCE | IDENTIFIER |
| --- | --- | --- |
| Rabbit polyclonal anti-Nanog | Abcam | Cat# ab80892,<br>RRID:<br>AB_2150114<br>Lot: GR40243-12 |
| Goat polyclonal anti-Sox2 | R&D Systems | Cat# AF2018,<br>RRID:<br>AB_355110 |
| Rabbit polyclonal anti-Stella | Santa Cruz | Cat# sc-67249,<br>RRID:<br>AB_1128372,<br>Lot: H1809 |
| Goat polyclonal anti-Stella | R&D Systems | Cat# AF2566,<br>RRID:<br>AB_2094147,<br>Lot:UWS010811<br>1 |
| Goat polyclonal anti-Oct4 | Santa Cruz | Cat# sc-8628,<br>RRID:<br>AB_653551,<br>Lot: F1815 |
| Rabbit polyclonal anti-Ap2γ | Santa Cruz | Cat# sc-8977,<br>RRID:<br>AB_2286995,<br>Lot: G1112 |
| Goat polyclonal anti-Brachyury (N-19) | Santa Cruz | Cat# sc-17743<br>RRID:<br>AB_634980<br>Lot: A1614 |
| Rat monoclonal anti-E-Cadherin | Sigma-Aldrich | Cat#U3254<br>RRID:<br>AB_477600<br>Lot: 085K4798 |
| Rabbit polyclonal anti-Eomes | Abcam | Cat#ab23345<br>RRID:<br>AB_778267<br>Lot:GR306193-1 |
| Rabbit monoclonal anti-phospho Smad1/5/9 | Cell Signaling Technology | Cat# 13820<br>RRID:<br>AB_2493181<br>Lot: D5810 |
| Goat polyclonal anti-Otx2 | R&D Systems | Cat# AF1979,<br>RRID:<br>AB_2157172,<br>Lot:<br>KNO0615111 |
| Rat monoclonal anti-Blimp1 | Santa Cruz | Cat# sc-130917,<br>RRID:<br>AB_2169704,<br>Lot: G0913 |
| Rabbit polyclonal anti-GFP Alexa Fluor 488 | Invitrogen | Cat# A21311,<br>RRID:<br>AB_221477,<br>Lot: 1891008 |
| Chicken polyclonal anti-GFP | Abcam | Cat# Ab13970,<br>RRID:<br>AB_300798,<br>Lot: GR305729-1 |
| Goat anti-chicken Alexa Fluor 488 | Invitrogen | Cat# A11039,<br>RRID:<br>AB_142924, |
| Donkey anti-rabbit Alexa Fluor 594 | Invitrogen | Cat# A21207,<br>RRID:<br>AB_141637 |
| Donkey anti-goat Alexa Fluor 594 | Invitrogen | Cat# A11058,<br>RRID:<br>AB_142540 |
| Donkey anti-rabbit Alexa 488 | Molecular Probes | Cat# A21206,<br>RRID:<br>AB_141708 |
| Donkey anti-rat 594 | Molecular Probes | Cat# A21209<br>RRID:<br>AB_2535795 |
| Rabbit anti-rat IgG | Vector<br>Laboratories | Cat# AI-4001<br>RRID:<br>AB_2336209 |

**Supplemental Table S4.**
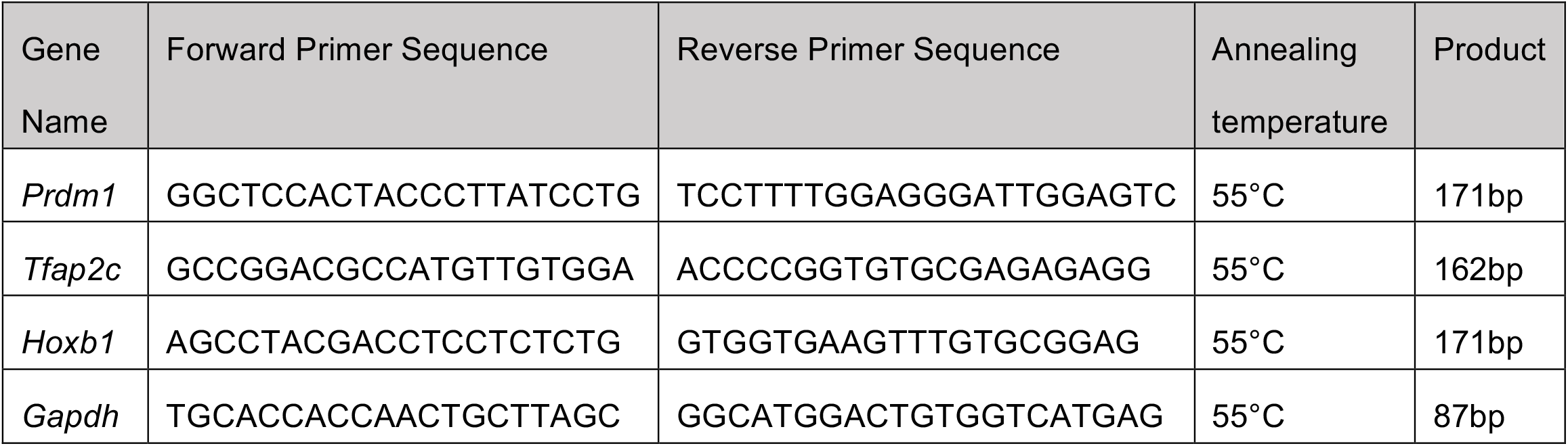

